# Epigenetic Regulation of Endothelial Extracellular Matrix Components is Critical for Murine Lung Development

**DOI:** 10.1101/2023.08.09.552718

**Authors:** Meng-Ling Wu, Kate Wheeler, Robert Silasi-Mansat, Florea Lupu, Courtney T. Griffin

**Affiliations:** Cardiovascular Biology Research Program, Oklahoma Medical Research Foundation, Oklahoma City, OK 73104, USA; Department of Cell Biology, University of Oklahoma Health Sciences Center, Oklahoma City, OK 73104, USA

## Abstract

**Background:** The chromatin remodeling enzymes BRG1 (brahma-related gene 1) and CHD4 (chromodomain helicase DNA binding protein 4) independently regulate transcription of genes critical for vascular development, but their coordinated impact on vessels in late-stage embryos has not been explored.

**Methods:** In this study we genetically deleted endothelial *Brg1* and *Chd4* in mixed background mice (*Brg1^fl/fl^;Chd4^fl/fl^;VE-Cadherin-Cre^+^*), and littermates that were negative for Cre recombinase were used as controls. Perinatal lung tissue was analyzed by immunostaining, immunoblots, and flow cytometry. Quantitative reverse transcription PCR was used to determine gene expression, and chromatin immunoprecipitation revealed gene targets of BRG1 and CHD4 in cultured endothelial cells (ECs).

**Results:** We found that *Brg1/Chd4* double mutants died soon after birth with small and compact lungs. Despite having normal cellular composition, distal air sacs of the mutant lungs displayed diminished ECM (extracellular matrix) components and TGFβ (transforming growth factor beta) signaling, which typically promotes matrix synthesis. Transcripts for collagen- and elastin-related genes and the TGFβ ligand *Tgfb1* were decreased in mutant lung ECs, but genetic deletion of endothelial *Tgfb1* failed to recapitulate the small lungs and ECM defects seen in *Brg1/Chd4* mutants. We instead found several ECM genes to be direct targets of BRG1 and CHD4 in cultured ECs.

**Conclusions:** Collectively, our data highlight essential roles for ECs in promoting ECM deposition at late stages of embryonic lung development. Moreover, this endothelial ECM production is epigenetically regulated.

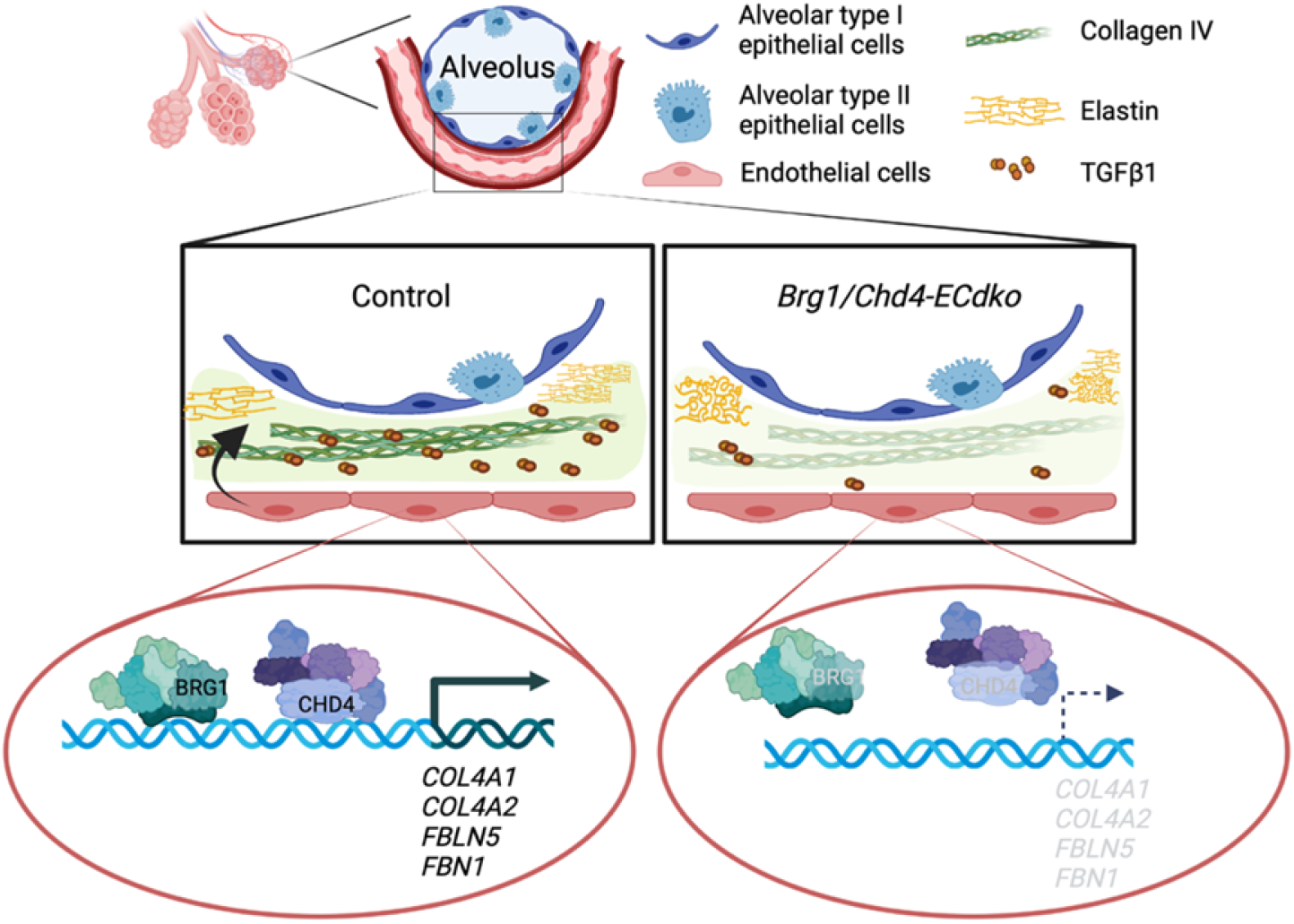

**HIGHLIGHTS:**

- Genetic deletion of the chromatin remodeling enzymes BRG1 and CHD4 in endothelial cells of late-stage mouse embryos (*Brg1/Chd4-ECdko*) results in small and compact lungs at birth.
- Mutant embryos display reduced collagen IV deposition, dysregulated elastin fibers, and diminished TGFβ1 in the distal air sacs.
- Our combined in vitro and in vivo analyses indicate that BRG1 and CHD4 epigenetically regulate collagen IV- and elastin-related gene expression in embryonic ECs to promote proper lung development.

## INTRODUCTION

ATP-dependent chromatin-remodeling complexes can epigenetically modulate gene expression by transiently disrupting histone-DNA contacts to change the density and spacing of nucleosomes in gene regulatory regions and allow access for transcriptional regulators^1^. Our lab previously exploited endothelial-specific deletion of catalytic ATPases within chromatin-remodeling complexes to identify genes that are required for embryonic vascular development. Chromodomain helicase DNA-binding 4 (CHD4) is an ATPase within the nucleosome remodeling deacetylase (NuRD) complex. We found that endothelial CHD4 regulates plasmin activity by suppressing excessive transcription of urokinase plasminogen activator (uPA) and its receptor (uPAR) to maintain vascular homeostasis during embryonic development^2,3^. We also investigated the ATPase brahma-related gene 1 (BRG1), which belongs to the switch/sucrose nonfermentable (SWI/SNF) complex. Embryonic deletion of *Brg1* reveals various roles for this enzyme in yolk sac vascular patterning^4,5^, venous identity^6^, capillary integrity^7^, and omental vascular homeostasis^8^.

Different chromatin-remodeling complexes can simultaneously act upon the same gene regulatory region to modulate expression of a target gene or pathway^9,10^. ChIP-Seq experiments indicate high co-occupancy of BRG1 and CHD4 across the genome of mouse mammary epithelial cells^9^. Moreover, BRG1 and CHD4 regulate *Tp53* and *Cdkn1a* gene expression cooperatively in P19 embryonic carcinoma cells^11^ but work antagonistically in regulating LPS-induced inflammatory genes in J774 macrophages^12^. Our lab previously reported an antagonistic role for BRG1 and CHD4 in regulating the Wnt signaling pathway in yolk sac endothelial cells (ECs) during early vascular development^5^, but the simultaneous regulation of genes or signaling pathways by these chromatin-remodeling enzymes in later stages of blood vessel development has not yet been explored.

In the current study, we addressed this gap in knowledge by deleting *Brg1* and *Chd4* with a constitutively active *VE-Cadherin-Cre* transgenic line that is fully penetrant in ECs by embryonic day 14.5 (E14.5) ^13^. We found that mutants deficient in both endothelial *Brg1* and *Chd4* displayed small and compact lungs around birth. Moreover, these lung defects correlated with a deficit of extracellular matrix (ECM) components. ECM is recognized for promoting lung development and homeostasis^14–16^, and fibroblasts/myofibroblasts are typically recognized as the major sources of pulmonary ECM^15,17–19^. Our data now indicate that ECs are also involved in ECM modulation during lung development and that BRG1 and CHD4 epigenetically regulate multiple ECM genes in ECs.

## MATERIALS AND METHODS

Extended materials and methods can be found in supplementary materials, and detailed information for resources used in the study is listed in the Major Resources Table.

### Mouse lines

*Brg1-floxed* mice^20^, *Chd4-floxed* mice^21^, *VE-Cadherin-Cre* transgenic mice^13^, *Tgfb1- floxed* mice^22^, and *Cdh5(PAC)-CreERT2* mice^23^ were genotyped as described^5,22,24,25^. *Brg1^fl/fl^;Chd4^fl/fl^*mice were bred to *Brg1^fl/fl^;Chd4^fl/+^;VE-Cadherin-Cre* to generate *Brg1/Chd4-ECdko* mutants and were maintained on a mixed genetic background. *Tgfb1^fl/fl^* mice were bred to *Tgfb1^fl/fl^;VE-Cadherin-Cre* mice to generate *Tgfb1-ECko* embryos and were maintained on a C57BL/6J background. *Chd4^fl/fl^* mice were bred to *Chd4^fl/fl^;Cdh5(PAC)-CreERT2* mice to generated inducible *Chd4^fl/fl^;Cdh5(PAC)- CreERT2* mutants. Mice bred for timed mating purposes were between eight weeks and one year of age. Noon on the day of vaginal plug detection was designated as E0.5. Littermates that were negative for Cre recombinase were used as controls to minimize potential genetic background effects. Experimental animals were selected based on genotype; small numbers of experimental (i.e., mutant) animals within each litter excluded a need for randomization of this category. Mutant animals were excluded from the study if their tissues did not show evidence of gene excision by PCR analysis. Data from male and female embryos were pooled into one group for analysis since no gender effects were observed. All mice were housed in the Oklahoma Medical Research Foundation (OMRF), which is an American Association for Accreditation of Laboratory Animal Care (AAALAC) accredited facility. All animal experiments were conducted under the approval of the OMRF Institutional Animal Care and Use Committee (IACUC).

### Lung weight measurement

Lungs were harvested from control and mutant pups 1 hour after cesarean section. Body weights and wet lung weights were determined immediately after isolation. Dry lung weight was determined after incubating each tissue at 60°C until the weight became consistent.

### Lung DNA content measurement

Total DNA content was isolated from left lungs (E16.5) or right lungs (E18.5) of control and mutants using TRIzol according to the manufacturer’s instructions. DNA concentration of samples was determined by NanoPhotometer C40-GO (IMPLEN).

### Human umbilical vein endothelial cell (HUVEC) culture and siRNA transfection

HUVECs (ATCC) were grown in EBM-2 media (Lonza) supplemented with EGM-2 SingleQuots (Lonza) and Antibiotic-Antimycotic (Gibco) according to the manufacturer’s recommendation. Cells were maintained in a 37°C incubator with a humidified atmosphere of 5% CO_2_. Only HUVECs below passage 8 were used for experiments in this study. For siRNA-mediated gene knockdown, HUVECs were plated in 6-well plates one day before transfection with BRG1-, CHD4-, or non-specific control siRNA oligos (Thermo Fisher Scientific, 100 pmole) using Lipofectamine RNAiMAX Transfection Reagent (Thermo Fisher Scientific) according to the manufacturer’s protocol. RNA was extracted for gene expression analysis 48 hours after transfection.

### Statistics

Prism 9.0 software (GraphPad) was used for all statistical assessments. Details for specific statistical analyses can be found in the figure legends. Tests for normality and equal variance were used to determine the appropriate parametric or non-parametric statistical model. In general, the comparison between two groups was assessed by unpaired parametric two-tailed t-tests. Comparison of multiple means was made using repeated-measures ANOVA with multiple comparisons between individual group means unless specified. Statistically analyzed data are presented as mean (±SD), associated numbers indicate adjusted p values, and dots on the bar graphs indicate biological replicates.

## RESULTS

### Deletion of endothelial *Brg1* and *Chd4* causes lung dysplasia

We previously deleted floxed alleles of either *Chd4* or *Brg1* with the *VE-Cadherin-Cre* recombinase and reported that *Chd4* mutants die with liver hemorrhage by E16.5, while *Brg1* mutants survive past birth without obvious vascular phenotypes^3,24^. To delete *Brg1* and *Chd4* simultaneously in ECs around midgestation, we generated *Brg1^fl/fl^;Chd4^fl/fl^;VE-Cadherin-Cre^+^* (hereafter referred to as *Brg1/Chd4-ECdko*) mouse embryos. We first found that 12 out of 22 *Chd4^fllfl^;VE-Cadherin-Cre^+^*(*Chd4-ECko*) mutants died at E14.5, but no lethality was seen in *Brg1^fl/fl^;Chd4^fll+^;VE-Cadherin-Cre^+^*(*Brg1-ECko*) or *Brg1/Chd4-ECdko* mutants dissected from 18 litters (Table S1). These data demonstrate that simultaneous deletion of endothelial *Brg1* rescues the lethality seen in *Chd4-ECko* mutants at E14.5. Although embryo size and weight appeared normal between littermate controls and mutants at E18.5 (Fig. 1A and 1C), we then found that *Brg1/Chd4-ECdko* lungs were small compared to control lungs, despite having normal lobation (Fig. 1B). Concordantly, after normalizing dry lung weights with body weights in control and mutant embryos, lung weight/body weight ratios were decreased in *Brg1/Chd4-ECdko* mutants compared to control littermates (Fig. 1D).

**Fig. 1.**
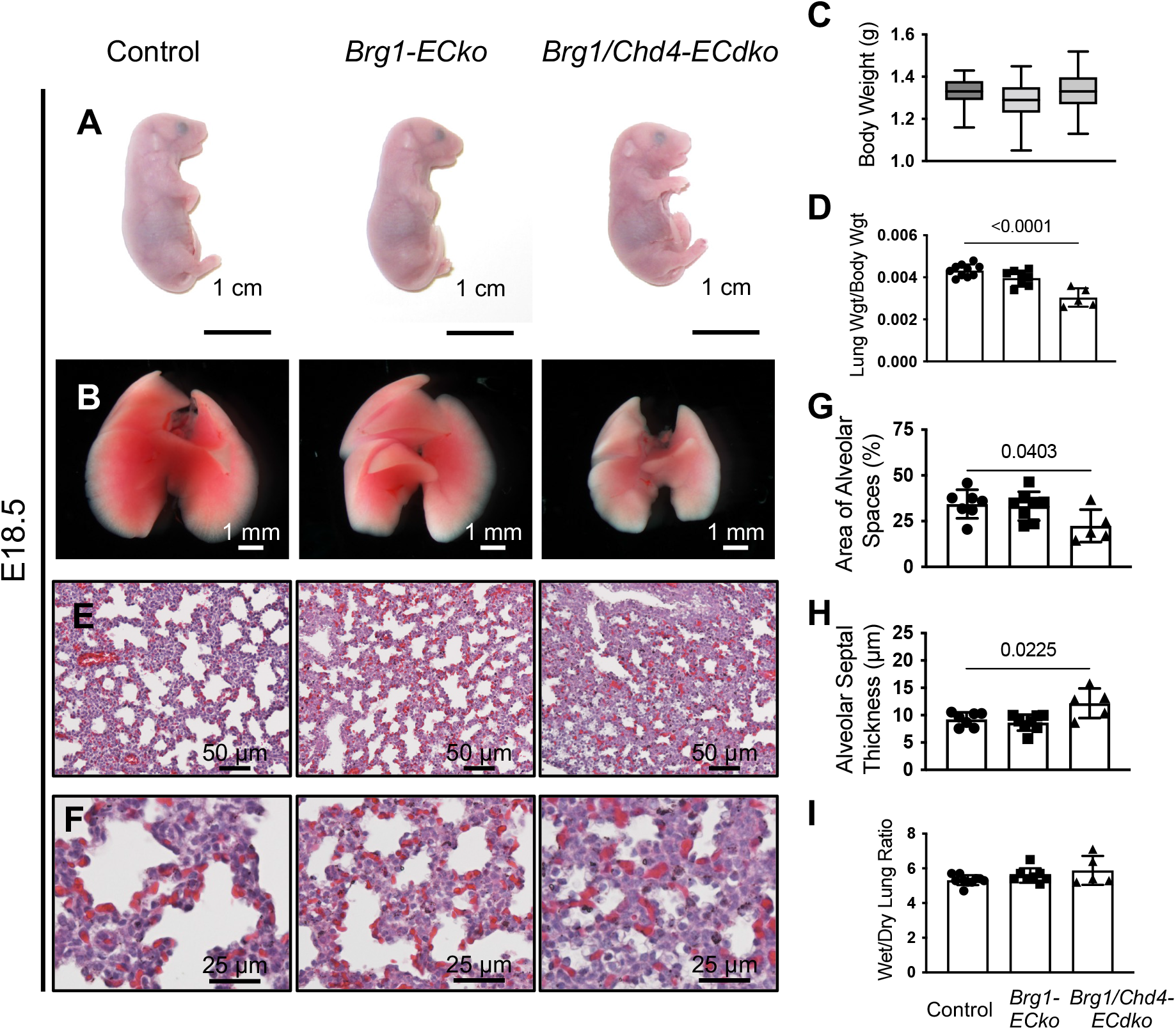
*Brg1/Chd4-ECdko* mutants have small and compact lungs. Representative pictures of littermate control, *Brg1-ECko*, and *Brg1/Chd4-ECdko* embryos (A) and lungs (B) at E18.5. (C) Body weight of E18.5 control and mutant embryos. Data are presented in box and whisker plots showing minimum to maximum (n ≥ 18 for each group). (D) Dry lung weight/body weight ratio at E18.5 (n ≥ 5). (E and F) Hematoxylin and eosin (H&E) staining of control and mutant lungs at E18.5. Quantification of mean alveolar space (G) and alveolar septal thickness (H) in E18.5 lungs from control and mutant embryos (n ≥ 5). (I) Wet/dry lung ratio after cesarean sections (C-sections) from E18.5 control and mutant embryos (n ≥ 5). Data are represented as mean (±D) and were analyzed with ordinary one-way ANOVA with Dunnett’s multiple comparisons post-test.

Notably, the developmental defects were restricted to *Brg1/Chd4-ECdko* lungs and did not occur in other organs such as the liver (Fig. S1A). Furthermore, histological analysis of H&E-stained lung sections revealed defects in the distal air sacs of the double-mutant lungs. *Brg1/Chd4-ECdko* lungs had significantly compacted alveolar spaces and thicker alveolar septa compared to control lungs at E18.5 (Fig. 1E-H), although lung size and histology were comparable between controls and mutants at E16.5 (Fig. S1B-C). Therefore, *Brg1/Chd4-ECdko* mutants developed small and compact lungs during the saccular stage of embryonic lung development.

Because we never saw *Brg1/Chd4-ECdko* mutants survive until weaning, we assessed whether *Brg1/Chd4-ECdko* mutants could survive after birth by performing cesarean sections (C-sections) on pregnant dams carrying E18.5 embryos. We then observed the pups for one hour after delivery. Fifty-five percent of *Brg1/Chd4-ECdko* pups died within 20 minutes with a cyanotic appearance (Fig. S1D-E). H&E staining of lung sections revealed compact lung phenotypes in both *Brg1-ECko* and *Brg1/Chd4-ECdko* mutants delivered by C-sections (Fig. S1F-G). Moreover, alveolar spaces were reduced, and alveolar septal thickness increased progressively in *Brg1-ECko* and *Brg1/Chd4-ECdko* lungs (Fig. S1H-I). Lymphatic function is required for lung inflation after birth, and loss of lymphatics causes perinatal lethality and compact lungs^26^. Since the *VE-Cadherin-Cre* recombinase is active in lymphatic ECs in addition to blood vessel ECs, we immunostained for the lymphatic EC marker VEGFR3 in E18.5 control and mutant lungs. *Brg1/Chd4-ECdko* mutants showed VEGFR3^+^ lymphatic vessels in their lungs (Fig. S1J) and their wet/dry lung weight ratio after C-sections delivery was normal (Fig. 1I). These data indicate that the compact lungs in *Brg1/Chd4-ECdko* mutant are not caused by lymphatic defects.

To exclude the possibility that the lung phenotypes we observed in *Brg1/Chd4-ECdko* mutants were induced solely by CHD4 deficiency, we generated tamoxifen-induced endothelial *Chd4* mutant embryos (*Chd4^fllfl^;Cdh5(PAC)-CreERT2^+^*). Tamoxifen was administered to pregnant dams from E12.5 to E14.5, and lung histology was analyzed at E18.5. We found that littermate control (*Chd4^fl/fl^*) and *Chd4^fl/fl^;Cdh5(PAC)-CreERT2^+^* mutants had comparable lung morphology (Fig. S1K). Additionally, alveolar spaces and septa looked normal in *Chd4* mutant lungs (Fig. S1L-M). These findings indicate that endothelial *Chd4* deletion is not sufficient to induce lung dysplasia comparable to that seen in E18.5 *Brg1/Chd4-ECdko* lungs.

### *Brg1/Chd4-ECdko* lungs have normal cell numbers and types

Various genetic mouse models show compact or dysplastic lung phenotypes like the ones described above in *Brg1/Chd4-ECdko* mutants^14,27–29^. Defective cell proliferation during lung development is one cause of pulmonary dysplasia^27^. To determine if the small lungs in E18.5 *Brg1/Chd4-ECdko* mutants resulted from insufficient cell proliferation during preceding developmental stages, we immunostained control and mutant lung sections at E16.5 and E17.5 for the proliferation marker phospho-histone H3 (pHH3). More pHH3^+^ cells were detected in E16.5 lungs than in E17.5 lungs across all genotypes, but proliferative cell numbers were comparable between genotypes at both E16.5 and E17.5 (Fig. 2A-B and S2A). The cell proliferation markers pHH3 and PCNA were also assessed by immunoblot in E16.5 lungs and were found to be comparable between genotypes (Fig. S2B). These results indicate that cell proliferation defects do not contribute to the small *Brg1/Chd4-ECdko* lungs seen at E18.5. Since increased cell apoptosis can also contribute to developmental lung hypoplasia^28^, we next assessed apoptotic cell numbers in control and mutant E17.5 lung sections using terminal deoxynucleotidyl transferase dUTP nick end labeling (TUNEL) staining. However, we did not observe any differences in apoptotic cell numbers between control and mutant genotypes (Fig. 2C), indicating that increased cell death likewise does not cause small *Brg1/Chd4-ECdko* lungs at E18.5.

**Figure 2.**
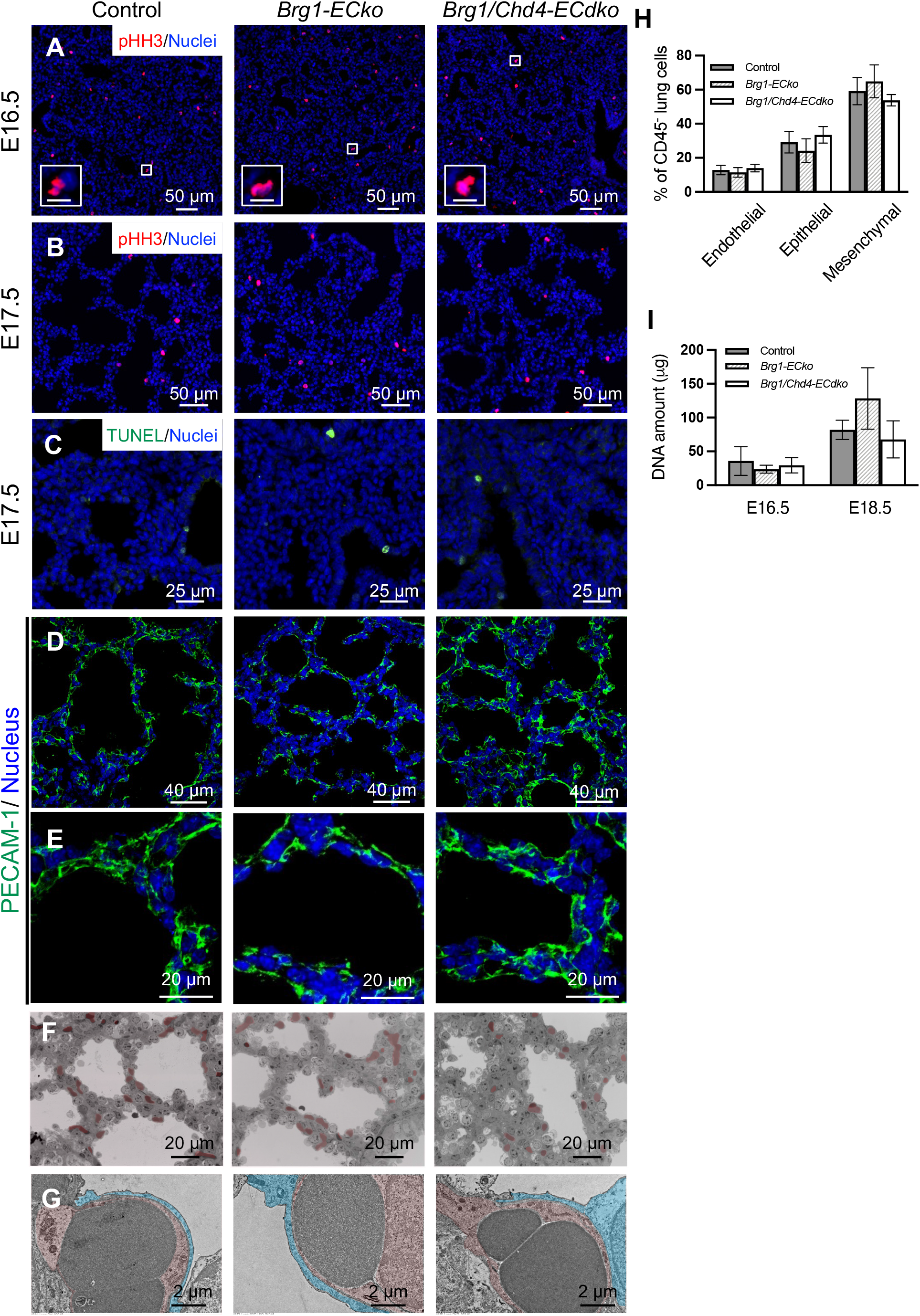
Cell numbers and cell types are normal in *Brg1/Chd4-ECdko* lungs. Proliferative cells were identified by phospho-histone H3 (pHH3, red) immunostaining of control, *Brg1-ECko*, and *Brg1/Chd4-ECdko* lung sections at E16.5 (A) and E17.5 (B). Nuclei were counterstained with Hoechst (blue). Boxes show magnified pHH3^+^ cells. Scale bars in boxes: 10 μm. (C) Apoptotic cells were identified by TUNEL staining (green) in E17.5 control and mutant lungs. (D&E) Vasculature in distal lungs was assessed by PECAM-1 (green) immunostaining of E18.5 control and mutant lungs. (F) Light microscopy of semithin sections from E18.5 control and mutant lungs. Red blood cells were pseudocolored in red to highlight capillaries in distal lungs. (G) Representative images of transmission electron microscopy (TEM) micrographs from E18.5 control and mutant lungs. Alveolar epithelial cells and ECs were pseudocolored in blue and red, respectively. (H) Cell populations within whole E18.5 lungs were determined by flow cytometry. Endothelial cells (PECAM-1^+^/CD45^-^), epithelial cells (EPCAM^+^/CD45^-^), and mesenchymal cells (PECAM-1^-^EPCAM^-^/CD45^-^) were determined after gating live CD45^-^ lung cells (n ≥ 3). (I) DNA content of control and mutant lungs at E16.5 and E18.5 (n ≥ 4). DNA amount was determined by measuring the A_260_ in a spectrophotometer. Data are represented as mean (±SD) and were analyzed with ordinary one-way ANOVA with Dunnett’s multiple comparisons post-test.

Aberrant cellular composition can also contribute to compact lung development. For example, vascular deficiency is associated with several hypoplastic lung models^28,29^. Therefore, we next assessed lung vascularity by immunostaining control and mutant lung sections for the EC marker PECAM-1, but we saw no paucity of PECAM-1^+^ vessels in *Brg1/Chd4-ECdko* mutants (Fig. 2D-E). Additionally, knowing that capillaries in distal lungs should closely abut alveolar spaces for efficient gas exchange, we analyzed semithin sections from E18.5 control and mutant lungs and found that capillaries in *Brg1/Chd4-ECdko* lungs showed normal apposition to the alveolar spaces (Fig 2F). Transmission electron microscopy (TEM) further revealed a normal interface between ECs and alveolar epithelial cells in *Brg1/Chd4-ECdko* lungs (Fig 2G).

We next sought to quantify the three major lung cell types (ECs, epithelial cells, and mesenchymal cells) by performing flow cytometry on E18.5 control and mutant lungs and found each cell population to be similar across genotypes (Fig. 2H). Since defects in alveolar epithelial cell differentiation are also reported in compact lungs^14,27^, we immunolabelled alveolar progenitor cells (SOX9^+^) and differentiated type II (pro-surfactant protein C^+^ = pro-SPC^+^) and type I (podoplanin^+^) alveolar epithelial cells in E18.5 control and mutant lungs, but once again we saw no differences between genotypes (Fig. S2C-E). Although pro-SPC^+^ cell numbers appeared to be increased in immunostained *Brg1/Chd4-ECdko* lung sections, no genotypic differences in type II cell numbers persisted after normalizing to nuclear cell counts (Fig. S2F). Finally, to assess the total cellular content in control and mutant lungs, we measured DNA content in E16.5 and E18.5 lungs and found similar levels of DNA across the genotypes (Fig. 2I). Altogether, these data indicate that deficiencies in cellular number and types do not contribute to small *Brg1/Chd4-ECdko* lungs.

### Collagen IV and elastin fibers are diminished in distal *Brg1/Chd4-ECdko* lungs

ECM is essential not only for the structural support of postnatal lungs but also for promoting lung development^15,30^. Since the cellular composition of embryonic *Brg1/Chd4-ECdko* lungs was normal, we next analyzed ECM components in *Brg1/Chd4-ECdko* lungs. Type IV collagen, the main component of the vascular basement membrane, is essential for the proper formation of pulmonary epithelium and vasculature and is likewise key for alveolar myofibroblast development^14^. We found collagen IV protein levels to be decreased in E18.5 *Brg1-ECko* and *Brg1/Chd4-ECdko* lungs by anti-collagen IV immunostaining of lung sections and by immunoblotting lung lysates (Fig. 3A and 3F). Elastin is another important pulmonary ECM component, which provides elastic recoil to the lung and is essential for its function^31^. Mice globally deficient for *Eln* (the gene name for the soluble elastin precursor tropoelastin) have emphysema-like lungs with dilated distal air sacs at birth^16^. Since collagen IV deficiency can lead to abnormal elastin fiber deposition^14^, we next visualized elastin fibers in control and mutant lung sections with Weigert’s resorcin fuchsin stain. We found elastin fibers to be fragmented and tortuous in distal air sacs of E18.5 *Brg1/Chd4-ECdko* lungs, while elastin fibers around arteries and bronchioles in proximal lungs were comparable at this stage (Fig. 3B-C and S3A). Moreover, immunostaining revealed tropoelastin to be diminished in *Brg1/Chd4-ECdko* lungs compared to controls (Fig. 3D), which was a trend also seen on immunoblots of *Brg1/Chd4-ECdko* lungs (Fig. 3G). To determine whether excessive matrix metalloproteinase (MMP) activity drove the diminished collagen and tropoelastin phenotypes in *Brg1/Chd4-ECdko* lungs, we next analyzed transcripts for MMPs that digest collagen IV or elastin and are known to be expressed in developing lungs^32^. Instead of seeing an increase in MMP transcripts as expected, we found *Mmp2* transcripts to be significantly reduced in *Brg1/Chd4-ECdko* lungs at E18.5, while *Mmp9*, *Mmp12*, and *Mmp13* transcripts were unaffected (Fig. S3B). In addition, MMP2 and MMP9 activities assessed by gelatin zymography were similar in control and mutant lungs (Fig. S3C).

**Figure 3.**
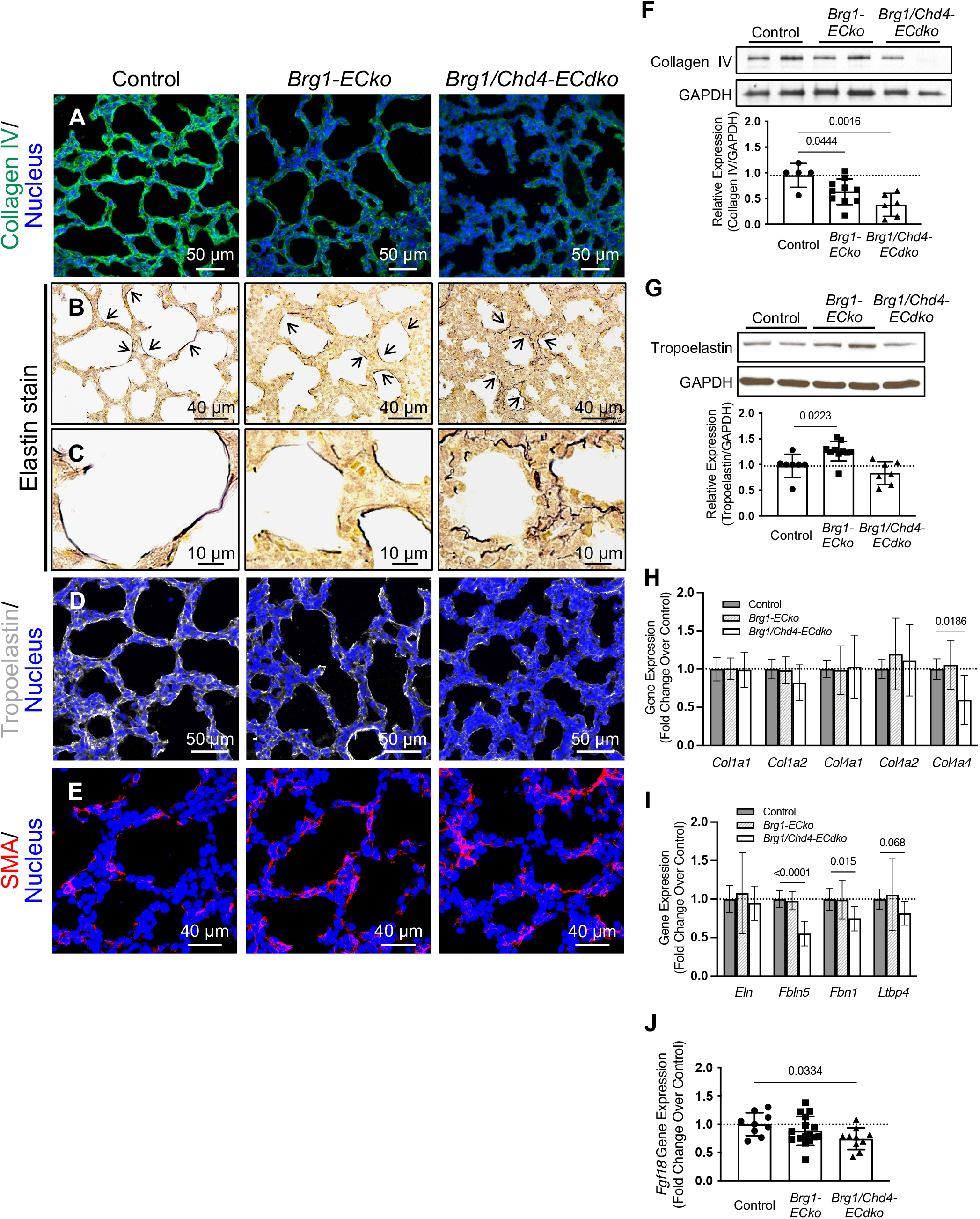
Collagen IV and elastin fibers are impaired in *Brg1/Chd4-ECdko* lungs. (A) Collagen IV expression was assessed by immunostaining of E18.5 control, *Brg1- ECko*, and *Brg1/Chd4-ECdko* lungs. Nuclei were counterstained with Hoechst (blue). (B&C) Elastin fibers (arrows) revealed by Weigert’s resorcin fuchsin staining are shown in black in E18.5 control and mutant lungs. Immunostaining for the soluble elastin precursor tropoelastin (D) and the myofibroblast marker α smooth muscle actin (SMA, E) are shown in E18.5 control and mutant lungs. Collagen IV (F) and tropoelastin (G) levels in E18.5 control and mutant lungs were assessed by immunoblot. Quantification below blot panels was determined by densitometry, and the dotted line shows the relative average expression of controls (n ≥ 5). Collagen- (H) and elastin-related (I) gene transcripts in E18.5 lungs were analyzed by qPCR (n ≥ 7). (J) *Fgf18* transcripts in E18.5 lungs were analyzed by qPCR (n ≥ 9). Data are represented as mean (±SD). Ordinary one-way ANOVA with Dunnett’s multiple comparisons post-test was used for (F-J). A Kruskal-Wallis test with Dunn’s multiple comparisons post-test was used for the *Ltbp4* gene due to a nonparametric distribution.

To determine if the reduced ECM protein levels seen in *Brg1/Chd4-ECdko* lungs were due to synthesis defects, we next examined collagen- and elastin-related gene transcripts in E18.5 control and mutant lungs by quantitative reverse transcription PCR (qPCR). We found that *Col4a4* transcripts were significantly reduced in *Brg1/Chd4- ECdko* lungs (Fig. 3H). Although *Eln* transcripts were not similarly reduced in *Brg1/Chd4-ECdko* lungs, we found that genes related to elastin fiber assembly (*Fbln5*, *Fbn1*, and *Ltbp4*) were decreased in *Brg1/Chd4-ECdko* lungs (Fig. 3I). These findings demonstrate that collagen IV synthesis is diminished, and that elastin fiber assembly is likely defective in *Brg1/Chd4-ECdko* lungs.

We next analyzed myofibroblasts using α smooth muscle actin (SMA) immunostaining since these cells are considered to be the primary source of ECM in developing lungs^15,17–19^. SMA^+^ myofibroblasts appeared comparable between *Brg1/Chd4-ECdko* and control lungs (Fig. 3E). We also analyzed transcripts for fibroblast growth factor 18 (*Fgf18*), since EC-derived retinoic acid (RA) signaling can promote elastin synthesis via mesenchymal FGF18 during alveolar development^33^. We found *Fgf18* transcripts were reduced in *Brg1/Chd4-ECdko* lungs (Fig. 3J), suggesting that mesenchyme-mediated elastin synthesis might be impacted in mutant lungs. However, although transcripts for one of the RA synthesis genes (retinaldehyde dehydrogenase 2; *Aldh1a2*) were reduced in *Brg1/Chd4-ECdko* lungs, none of the RA target gene transcripts we analyzed (*Egr1*, *Foxa1*, and *Pbx1*) were reduced in mutant lungs (Fig S3D-E). Therefore, ECM deficits in *Brg1/Chd4-ECdko* lungs are unlikely to be due to a deficit of myofibroblasts or to defective paracrine RA signaling between ECs and mesenchymal cells.

### TGFβ1 and its downstream signaling signature are diminished in *Brg1/Chd4- ECdko* lungs

TGFβ is another growth factor that is widely recognized for its ability to promote ECM production and to contribute to lung development^34,35^. Global deletion of genes encoding TGFβ2 or TGFβ3 results in perinatal defects in pulmonary distal air sacs^36,37^. *Tgfb2*-null mice have collapsed distal airways^37^, and *Tgfb3*-null mice show alveolar hypoplasia and mesenchymal thickening like we observed in *Brg1/Chd4-ECdko* lungs^36^. To determine if TGFβ signaling activity was impaired in *Brg1/Chd4-ECdko* lungs, we examined phosphorylation of the TGFβ effector molecule SMAD2 (p-SMAD2) through immunostaining and immunoblot at E18.5. We found p-SMAD2 levels to be significantly reduced in *Brg1/Chd4-ECdko* lungs compared to control lungs, and the reduction was pronounced in distal alveolae (Fig. 4A-C). Additionally, transcripts for the TGFβ target gene *Ctgf* were decreased in *Brg1/Chd4-ECdko* lungs at E18.5 (Fig. 4D). To determine whether specific TGFβ family members were diminished in *Brg1/Chd4-ECdko* lungs, we performed qPCR analysis and found that *Tgfb1* was significantly reduced, while *Tgfb2* transcripts trended downward (Fig. 4E). We next assessed *Tgfb1* expression in a publicly available single cell (sc)RNA-seq dataset (LungGENS) and found that ECs are among the cell types expressing *Tgfb1* in E18.5 lungs (Fig. 4F) ^38–40^. Moreover, among all *Tgfb1*-expressing cells in E18.5 lungs, 42.15% of them are ECs^38–40^. To determine if *Tgfb1* transcripts were downregulated in *Brg1/Chd4-ECdko* lung ECs, we isolated PECAM-1^+^ pulmonary ECs (PECs) from E18.5 control and mutant lungs and analyzed transcripts by qPCR. Transcripts of *Brg1* and *Chd4* were significantly reduced in PECs isolated from mutant lungs (Fig. S4A). *Tgfb1* transcripts were likewise significantly reduced in *Brg1/Chd4-ECdko* PECs (Fig. 4G). Therefore, we questioned whether reduced endothelial *Tgfb1* might contribute to the ECM deficits and compact lungs seen in E18.5 *Brg1/Chd4-ECdko* mutants.

**Figure 4.**
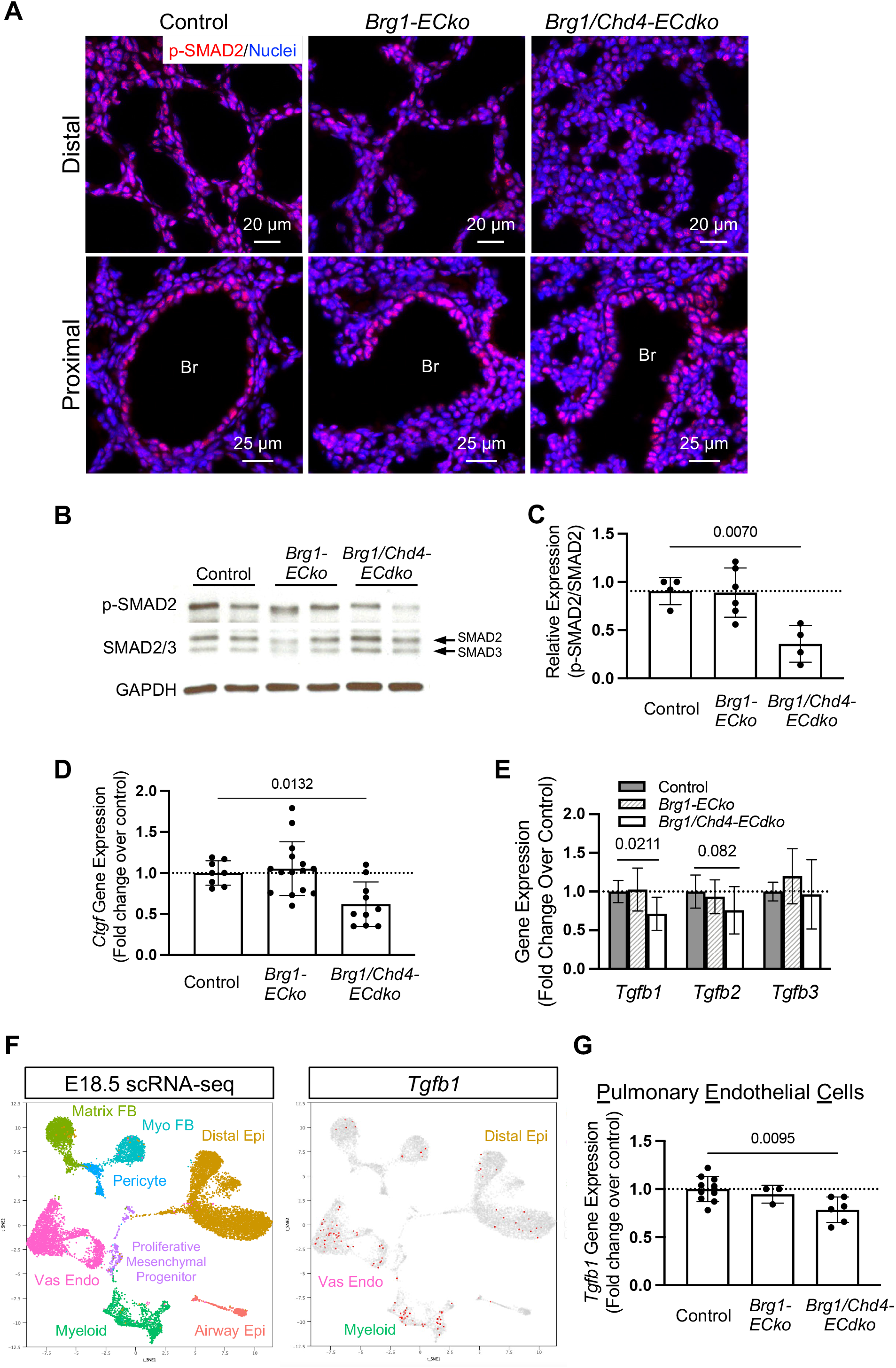
TGFβ1 and downstream signaling are reduced in *Brg1/Chd4-ECdko* lungs. (A) Phospho-SMAD2 (p-SMAD2) immunostaining of distal and proximal lungs in control, *Brg1-ECko*, and *Brg1/Chd4-ECdko* embryos at E18.5. Br: bronchioles. (B) pSMAD2 expression was assessed by immunoblot in E18.5 control and mutant lungs. (C) Relative pSMAD2 expression normalized by SMAD2 expression was determined by densitometry, and the dotted line shows the relative expression of controls (n≥ 4). (D) Transcripts for the TGFβ target gene *Ctgf* were assessed by qPCR in E18.5 control and mutant lungs (n≥ 8). (E) Gene transcripts of TGFβ family members *Tgfb1*, *Tgfb2*, and *Tgfb3* were assessed by qPCR in E18.5 lungs (n≥ 8). (F) t-SNE plots of cell clusters (left) and *Tgfb1* expression (right) within E18.5 lungs; data were extracted from the online LungGENS scRNA-seq database (https://research.cchmc.org/pbge/lunggens/default.html). Vas Endo: vascular ECs; Matrix FB: matrix fibroblasts; Myo FB: myofibroblasts; Distal Epi: distal epithelial cells; Airway Epi: airway epithelial cells. (G) *Tgfb1* transcripts in pulmonary ECs isolated from control and mutant embryos at E18.5 were assessed by qPCR (n≥ 3). Data are represented as mean (±SD) and were analyzed with ordinary one-way ANOVA with Dunnett’s multiple comparisons post-test.

### Deleting endothelial *Tgfb1* does not cause compact lungs and ECM loss during lung development

*Tgfb1* deletion from mesenchymal cells causes lung hypoplasia^28^, but the impact of endothelial *Tgfb1* deletion on lung development is unknown. To determine whether endothelial *Tgfb1* contributed to lung development by impacting collagen IV and elastin production during the late stages of embryonic development, we generated endothelial *Tgfb1* knockout embryos (*Tgfb1^fl/fl^;VE-Cadherin-Cre^+^*, hereafter referred to as *Tgfb1- ECko*). H&E staining of lung sections from E18.5 littermate control and *Tgfb1-ECko* mutants revealed similar lung histology, even at distal air sacs (Fig. 5A-B). Neither alveolar space area nor alveolar septal thickness was affected in *Tgfb1-ECko* lungs (Fig. 5C-D). To assess ECM components, we examined gene transcripts for collagen IV subunits and for the elastin assembly genes *Fbln5*, *Fbn1*, and *Ltbp4* in control and *Tgfb1-ECko* lungs. Although *Tgfb1* transcripts were significantly reduced in *Tgfb1-ECko* lungs, none of the ECM-related genes we tested were affected at E18.5 (Fig. 5E). We also analyzed collagen IV and tropoelastin protein levels by immunoblot and found their expression to be comparable in lungs from littermate control and *Tgfb1-ECko* mutants (Fig. 5F and S5A-B). Altogether, *Tgfb1-ECko* lungs did not phenocopy the compact morphology and ECM deficits we saw in E18.5 *Brg1/Chd4-ECdko* lungs.

**Figure 5.**
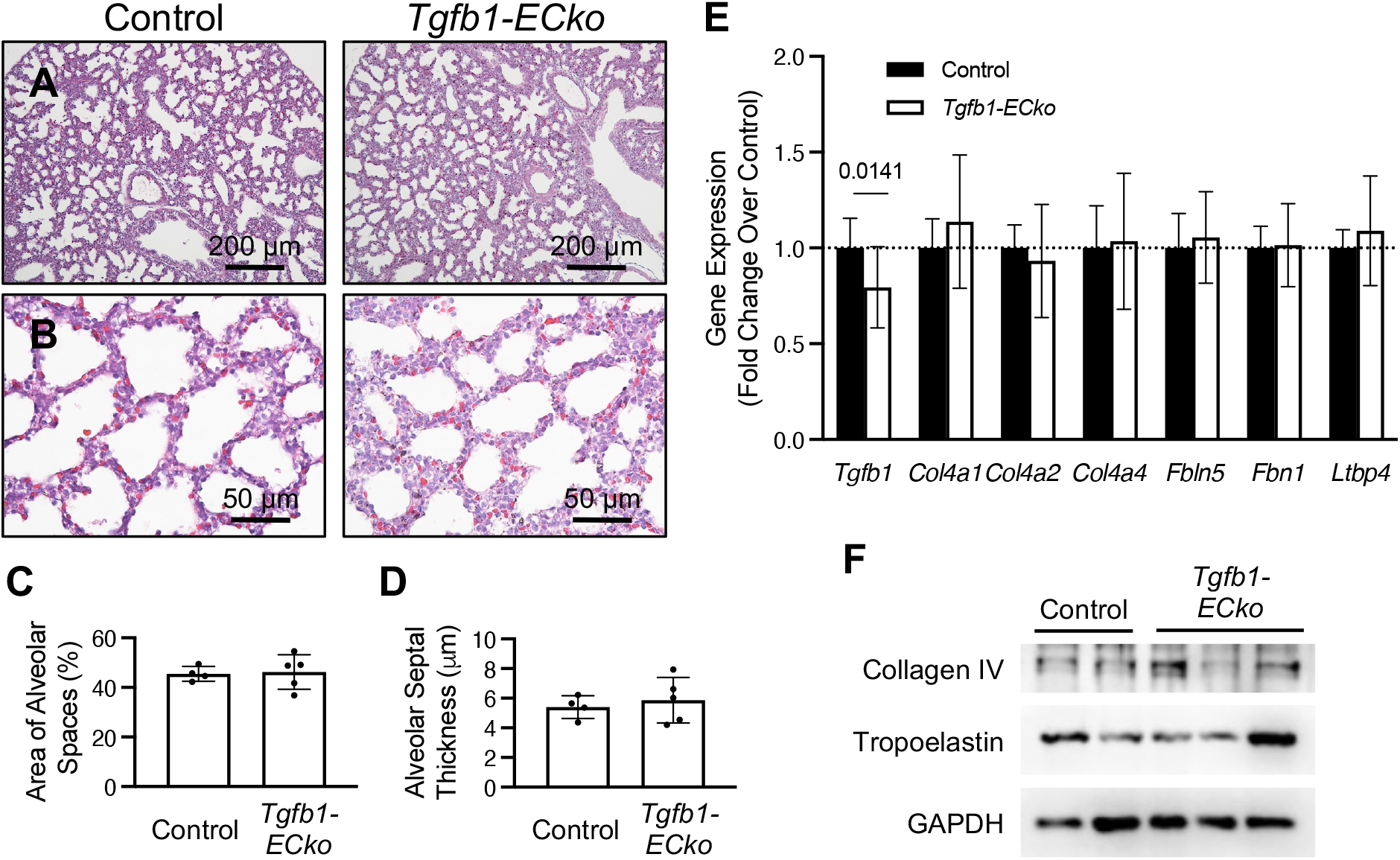
*Tgfb1-ECko* mice do not recapitulate *Brg1/Chd4-ECdko* lung phenotypes. (A&B) H&E staining of littermate control and *Tgfb1-ECko* lung sections at E18.5. Quantification of mean alveolar space (C) and alveolar septal thickness (D) in E18.5 control and *Tgfb1-ECko* lungs (n ≥ 4). (E) Transcripts of *Tgfb1* and ECM-related genes were assessed by qPCR in E18.5 control and *Tgfb1-ECko* lungs (n ≥ 6). (F) Collagen IV and tropoelastin expression in E18.5 control and *Tgfb1-ECko* lungs were assessed by immunoblot. Data are represented as mean (±SD) and were analyzed with unpaired t-tests.

### BRG1 and CHD4 directly regulate transcription of collagen IV subunits and elastin assembly genes in ECs

Since endothelial TGFβ1 did not act as an intermediate factor to drive synthesis of the ECM-related transcripts we analyzed during lung development, we next hypothesized ECs might be a direct source of ECM synthesis during lung development. This hypothesis is informed by the knowledge that ECs are the major source of elastin for the internal elastic lamina in muscular and resistance arteries^41^. We found through TEM analysis that ECs shared close contact with elastin and collagen fibers in subendothelial spaces of E18.5 distal lungs (Fig. 6 A-B). We then assessed whether BRG1 and CHD4 could directly regulate collagen IV and elastogenesis gene transcripts in ECs. Among the six known collagen IV alpha chain subunits, *COL4A1*, *COL4A2*, *COL4A5*, and *COL4A6* are detected in human pulmonary microvascular ECs^42^. Additionally, scRNA-seq data accessed through LungGENS indicated that ECs express *Col4a1* and *Col4a2* in E18.5 lungs (Fig. S6A) ^38–40^. Using qPCR, we found transcripts of *Col4a1* and *Col4a2* to be significantly downregulated in *Brg1/Chd4-ECdko* PECs isolated from E18.5 embryos (Fig. 6C). We likewise found transcripts of elastogenesis-related genes (*Eln*, *Fbln5*, *Fbn1*, and *Ltbp4*) to be significantly reduced in *Brg1/Chd4-ECdko* PECs (Fig 6D). Importantly, those same ECM-related gene transcripts were not impaired in non-EC cellular fractions of E18.5 *Brg1/Chd4-ECdko* lungs (Fig. S6B-C). These results indicated that synthesis of EC-derived ECM components was compromised in *Brg1/Chd4-ECdko* lungs. To assess whether BRG1 and CHD4 likewise impacted transcription of ECM components in cultured ECs, we knocked down CHD4 and/or BRG1 in human umbilical vein endothelial cells (HUVECs). We found *COL4A1* and *FBLN5* to be significantly downregulated and *FBN1* to be significantly upregulated after the simultaneous knockdown of both chromatin remodeling enzymes in these primary human ECs (Fig. 6E).

**Figure 6.**
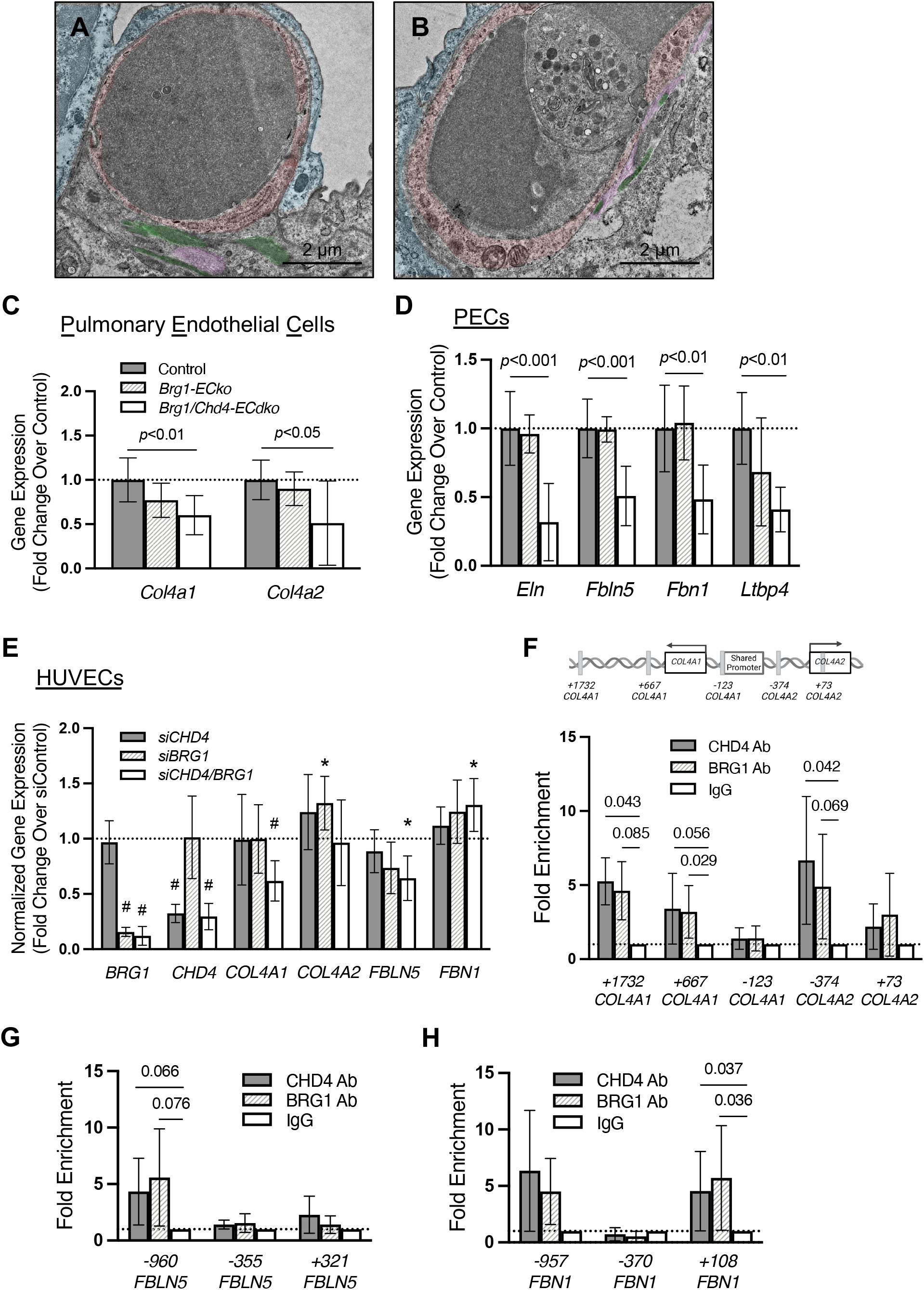
BRG1 and CHD4 directly regulate transcription of ECM components in ECs. Representative images of transmission electron microscopy (TEM) micrographs from E18.5 control (A) and *Brg1/Chd4-ECdko* (B) lungs. Alveolar epithelial cells (blue) and ECs (red) were pseudocolored. Collagen and elastin were pseudocolored in pink and green, respectively. (C) *Col4a1* and *Col4a2* transcripts were assessed by qPCR from isolated control, *Brg1-ECko*, and *Brg1/Chd4-ECdko* pulmonary ECs (PECs) at E18.5 (n≥ 3). (D) Transcripts of *Eln* and genes related to elastin fiber assembly (*Fbln5*, *Fbn1*, and *Ltbp4*) were assessed by qPCR in E18.5 PECs (n≥ 3). The dotted line shows the relative average expression of controls. (E) *CHD4* and/or *BRG1* were knocked down in HUVECs, and gene transcripts for *BRG1*, *CHD4*, *COL4A1*, *COL4A2*, *FBLN5*, and *FBN1* were assessed by qPCR 48 hours after the knockdown (n≥ 4). The dotted line shows the relative average expression of siControls. \**P*<0.05; ^#^*P*<0.01. (F) Upper panel: schematic diagram of the *COL4A1* and *COL4A2* gene loci, including regions tested for BRG1 and CHD4 enrichment. Lower panel: BRG1 and CHD4 enrichment at the *COL4A1* and *COL4A2* gene loci was assessed by chromatin immunoprecipitation (ChIP)-qPCR assays in HUVECs. DNA immunoprecipitated by antibodies against CHD4, BRG1, or IgG was amplified by qPCR, and data were expressed as fold enrichment over the level of ChIP with a negative control IgG antibody (dotted line, n≥ 3). Numbers for each amplicon were assigned based on the average distance of the amplified region relative to its nearest transcription start site (TSS). ChIP-qPCR analysis of CHD4 and BRG1 enrichment at the *FBLN5* (G) and *FBN1* (H) gene loci in HUVECs (n≥ 3). Data are represented as mean (±SD). Ordinary one-way ANOVA with Dunnett’s multiple comparisons post-test was used for analysis in C and D; one-sample t and Wilcoxon tests were used for E-H.

To determine whether some of the ECM-related genes that we saw misregulated in *Brg1/Chd4-ECdko* PECs and in *BRG1*/*CHD4*-knockdown HUVECs were direct targets of BRG1 and CHD4, we assessed binding of CHD4 and BRG1 to those ECM-related gene loci. We first used a comparative sequence alignment program (www.DCODE.org) to identify conserved regions of the *COL4A1*, *COL4A2*, *FBLN5*, and *FBN1* promoters in various species (Fig. S6D-F). We next performed ChIP-qPCR in HUVECs using either CHD4 or BRG1 antibodies. Since *COL4A1* and *COL4A2* are paralogous genes that share a bidirectional promoter^43,44^, we initially assessed this shared region but found no enrichment for either CHD4 or BRG1 at 123 bp upstream (-123) of the *COL4A1* transcription start site (TSS). However, we did find robust enrichment of both enzymes 374 bp upstream (-374) of the *COL4A2* TSS (Fig. 6F). In addition, we found enrichment of both BRG1 and CHD4 at intronic regions within the *COL4A1* gene (+1732 and +667) that are known to impact its transcription^44^. Our ChIP-qPCR analysis of the *FBLN5* gene likewise revealed enrichment of CHD4 and BRG1 in a conserved region of the promoter located 960 bp upstream (-960) of the TSS (Fig. 6G). We also found both CHD4 and BRG1 to be significantly enriched at 108 bp downstream (+108) of the *FBN1* TSS (Fig. 6H). Altogether, our data indicate that BRG1 and CHD4 can directly and simultaneously regulate *COL4A1*, *COL4A2*, *FBLN5*, and *FBN1* genes in ECs.

## DISCUSSION

Although ECs have been recognized as sources of ECM in various organs, the critical importance of endothelial-derived ECM deposition during lung development has been underappreciated^41,45–47^. In the present study, our EC-specific *Brg1/Chd4* double mutant mouse embryos provided a unique platform for revealing ECM genes that require tight transcriptional regulation during lung development. Loss of endothelial *Brg1* and *Chd4* resulted in small and compact lungs at the saccular stage of pulmonary development. This small lung phenotype was associated with defective expression of ECM components like collagen IV and elastin fiber assembly genes rather than changes in the cellular composition of the mutant lungs. At first, we thought the ECM deficits might be secondary to the downregulated endothelial *Tgfb1* transcripts and TGFβ signaling that we also discovered in *Brg1/Chd4-ECdko* mutant lungs. However, in our hands endothelial *Tgfb1* deletion was insufficient to cause the small lung phenotype and ECM deficits that we observed in *Brg1/Chd4-ECdko* mutant lungs. This finding refocused our attention on ECs, and we discovered that BRG1 and CHD4 coordinate to regulate transcription of several endothelial ECM components directly. Therefore, this study demonstrates the essential role of epigenetically regulated endothelial ECM deposition at the saccular stage of lung development.

Fibroblasts and myofibroblasts have typically been viewed as the main sources of ECM in developing and adult lungs. However, various genetic mutants have hinted at additional sources of pulmonary ECM. For example, mutants with a deficit of smooth muscle actin-positive secondary crest myofibroblasts at distal air sacs of the neonatal lung have normal pulmonary elastin expression, suggesting that non-myofibroblasts contribute to perinatal elastin synthesis^48^. Furthermore, deletion of the growth factor gene *Vegfa* in alveolar epithelial cells disrupts the distal pulmonary capillary network and impairs elastin deposition from fibroblasts, which suggests an indirect connection between ECs and elastin deposition^33^. Concordantly, pulmonary ECs isolated from different species secrete collagen IV and elastin *in vitro*^42,49,50^. Therefore, our genetic evidence in this study that ECs are a critical source of ECM components in the developing lung provides concrete support for these previous observations.

Our initial suspicion that diminished TGFβ1 signaling in *Brg1/Chd4-ECdko* lungs might be responsible for reduced ECM production was logical, particularly because TGFβ signaling is a major stimulus for ECM synthesis in myofibroblasts and because *Tgfb1* transcripts are predominantly found in perinatal pulmonary ECs according to scRNA-Seq data assembled in LungGENS^38–40^. However, the impact of TGFβ1 on lung development is controversial. *Tgfb1* global deletion does not cause perinatal lung phenotypes^51^, but cell-specific *Tgfb1* deletion impairs different aspects of lung development^28,52^. Although we did not find endothelial *Tgfb1* deletion to be sufficient to cause compact lungs and ECM loss in this study, it is still possible that reduced TGFβ signaling synergistically contributes to the ECM synthesis defects seen in *Brg1/Chd4- ECdko* lungs.

We showed in this study that multiple ECM components were impaired in *Brg1/Chd4- ECdko* lungs, including COL4A1, COL4A2, and elastin fibers. All these components are known to be important contributors to lung development. Homozygous mutations of *Col4a1* and/or *Col4a2* result in embryonic lethality at midgestation^53,54^, while heterozygous mutations in *Col4a1* lead to lethal lung defects around birth^14^. These mutants display defects in alveolar epithelial cell differentiation, vascular and myofibroblast development, and blood-gas barrier formation. Mice with global deletion of *Eln* have emphysema-like lungs with dilated distal air sacs at birth, and no mutants survive beyond 3.5 days after delivery^16^. Mice with global deletion of the gene encoding the elastogenesis protein Fibulin 5 (*Fbln5*) have severe emphysema-like lung histology with disrupted distal airways and fragmented elastin fibers at P3^55^. However, despite this defective development of elastic fibers, the mutants survive for at least six months with normal elastase activity. Global deletion of another elastogenic component, *Fbn1*, results in postnatal lethality and anomalies in multiple organ systems, including the lungs^56,57^. *Fbn1^-/-^*lungs are small and compact with thickened alveolar walls, like those seen in *Brg1/Chd4-ECdko* mutants. Because BRG1 and CHD4 are not transcription factors but rather function in complexes that modestly modify transcription of multiple target genes, we do not believe that any one of the ECM genes listed above is transcriptionally reduced enough to account fully for the *Brg1/Chd4-ECdko* lung phenotypes. Instead, we suspect that the anomalies seen in *Brg1/Chd4-ECdko* lungs are caused by combinatorial transcriptional reduction of multiple ECM components and potentially by *Tgfb1* in pulmonary ECs. Importantly, the genetic approach used in our study is designed to reveal endothelial BRG1 and CHD4 target genes that require tight transcriptional regulation, and our results indicate that the dosage of endothelial ECM component expression is important for proper lung development.

Genetic background strains impact baseline transcriptional networks in lung tissues of adult mice^58^. Although we bred *Brg1/Chd4-ECdko* mutants on a mixed genetic background, we did observe that the mutant lung phenotypes we observed were influenced by the composition of the background strain. Most notably, we lost the compact lung phenotype when the *Brg1/Chd4-ECdko* background strain was enriched for C57BL/6J, and we regained the phenotype when mutants were outbred to the CD1 background. Interestingly, the midgestational lethality of *Tgfb1^-/-^*mice is background sensitive, and two modifier genes of this phenotype have been identified^59,60^. Similarly, the severity of intracerebral hemorrhage (ICH) induced by *Col4a1* deletion is subject to genetic background^46^. *Col4a1* mutants crossed to the CAST/EiJ strain show minor ICH, while mutants crossed to the C57BL/6J strain have more severe intracellular COL4A1 accumulation and bleeding. Ultimately, we do not know the underlying modifier gene(s) that impacted the lungs of *Brg1/Chd4-ECdko* mutants in our study, and we recognize that considerable work would be needed to identify them. The *Tgfb1-ECko* mutants we generated were predominantly bred on the C57BL/6J background, and we acknowledge that pulmonary defects might emerge if we outbred these mutants further. However, we have not yet observed any lung phenotypes in *Tgfb1-ECko* mutants even after three backcrosses to the CD1 background. Therefore, we maintain that the *Brg1/Chd4-ECdko* lung defects are likely due to a combination of multiple misregulated genes and particularly endothelial ECM components.

Altogether, this study adds to our growing appreciation of the importance of EC-derived ECM in pulmonary development. Moreover, our discovery that endothelial ECM genes require tight epigenetic regulation during lung development raises interesting questions about their contribution to lung diseases. Given that chromatin remodeling enzymes can reactivate developmental gene expression programs following a pathological challenge^61^, we question whether endothelial ECM synthesis becomes epigenetically stimulated in the context of postnatal lung diseases. For this reason, additional focus on pulmonary endothelial ECM components and their regulation is warranted.

## ACKNOWLEDGMENTS

We thank Jun Xie for assistance with mouse colony maintenance. We also thank Siqi Gao, Jenna Chan, and Srija Nuguri for technical help and Griffin lab members for thoughtful discussions. We thank Jasimuddin Ahamed for *Tgfb1*-floxed mice, Katia Georgopoulos for *Chd4*-floxed mice, Pierre Chambon for *Brg1*-floxed mice, Lijun Xia for the podoplanin antibody, the OMRF Flow Cytometry Core Facility for assistance with flow cytometry, the OMRF Imaging Core Facility for paraffin block embedding, and we acknowledge primer sequences from PrimerBank (Massachusetts General Hospital and Harvard Medical School).

## AUTHOR CONTRIBUTIONS

Conceptualization, M.W. and C.T.G.; Validation, M.W., K.W., R.S., F.L. and C.T.G.; Formal analysis, M.W.; Investigation, M.W. and K.W.; Resources, M.W., F.L. and C.T.G.; Writing—Original Draft, M.W.; Writing—Review & Editing, M.W. and C.T.G.; Visualization, M.W. and C.T.G.; Supervision, C.T.G.; Funding Acquisition, M.W., and C.T.G.

## Supplemental Material

Supplemental Methods Supplemental Figures S1-S6 Supplemental Tables S1-S2 Major Resources Table

